# Structure-based screening of drug candidates targeting the SARS-CoV-2 envelope protein

**DOI:** 10.1101/2021.08.25.457645

**Authors:** Xin Xia, Yuwei Zhang, Songling Li, Hengwei Lin, Zhiqiang Yan

## Abstract

The COVID-19 (coronavirus disease 2019) pandemic is caused by SARS-CoV-2 (severe acute respiratory syndrome coronavirus 2). SARS-CoV-2 produces a small hydrophobic envelope (E) protein which shares high homology with SARS-CoV E protein. By patch-clamp recording, the E protein is demonstrated to be a cation-selective ion channel. Furthermore, the SARS-CoV-2 E protein can be blocked by a SARS-CoV E protein inhibitor hexamethylene amiloride. Using structural model and virtual screening, another E protein inhibitor AZD5153 is discovered. AZD5153 is a bromodomain protein 4 inhibitor against hematologic malignancies in clinical trial. The E protein amino acids Phe23 and Val29 are key determinants for AZD5153 sensitivity. This study provides two promising lead compounds and a functional assay of SARS-CoV-2 E protein for the future drug candidate discovery.

## INTRODUCTION

Till April 8, 2021, more than 133 million cases of COVID-19 and over 2.9 million deaths have been reported. There is currently no specific drug to treat COVID-19. It is an imperative that we develop drugs targeting SARS-CoV-2 for the treatment of COVID-19 infections.

Multiple highly pathogenic viruses encode small integral membrane proteins with ion channel activity (1). Ion channels of virus are critical for viral pathogenicity and can serve as ideal drug targets, for instance, amantadine, an anti-influenza A virus drug, is an ion channel antagonist (1). All coronaviruses including SARS-CoV-2 encode a small hydrophobic envelope (E) protein. Previous studies have shown that the Severe Acute Respiratory Syndrome Coronavirus (SARS-CoV) E protein forms ion channel and it is crucial for viral virulence and pathogenesis, including the formation of edema, the major determinant of pulmonary damage and acute respiratory distress syndrome (ARDS) (2-4).The E proteins of SARS-CoV and SARS-CoV-2 share high homology (94.7%, Figure. 1A). Given the high sequence identity, it is very likely that the E protein of SARS-CoV-2 might be crucial for viral pathogenicity and function as an ion channel.

**Figure 1.**
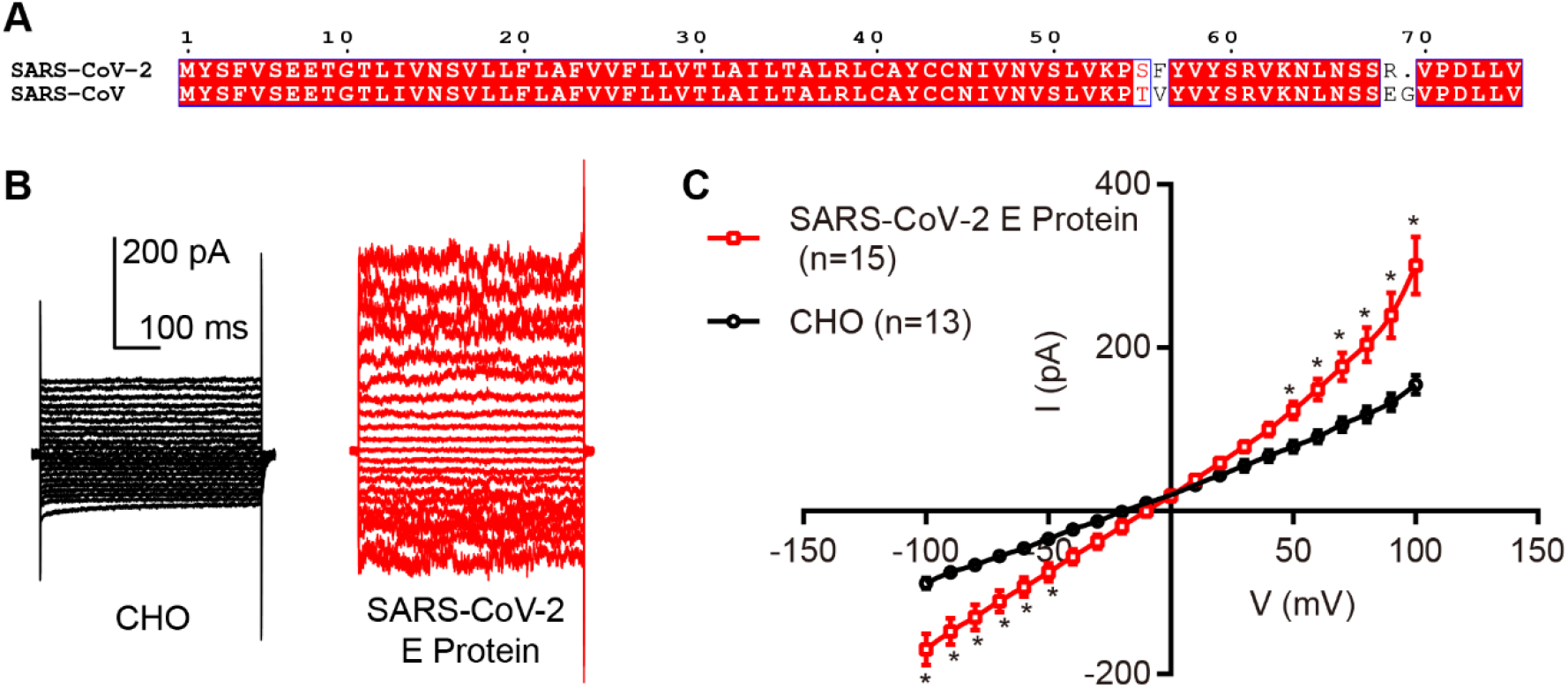
SARS-CoV-2 E protein forms ion channel. **(A)** The amino acid sequence alignment of SARS-CoV and SARS-CoV-2 E proteins (NCBI Reference Sequence: SARS-CoV E: NP_828854.1; SARS-CoV-2 E: YP_009724392.1). Strictly conserved residues are highlighted with a red box. The sequence alignment figure was generated by Clustal Omega and ESPript 3.0. **(B)** Sample traces of untransfected CHO cells, cells transfected with vector, cells expressing SARS-CoV-2 E protein. The cells were held at 0 mV. Step voltages ranged from - 100 to 100 mV with 10 mV increments. **(C)** Whole-cell current-voltage relationships of the indicated cells. The SARS-CoV-2 E gene showed moderate inward and relatively larger outward currents under voltage stimulation. The empty CHO cells showed differences compared with SARS-CoV-2 E protein transfected cells. Data are mean ± SEM. Statistical analysis used T-test; *, p<0.05.

## RESULTS

### SARS-CoV-2 E protein forms cation-selective ion channel

To test if E protein of SARS-CoV-2 is indeed an ion channel, the full sequence was synthesized and expressed in CHO cells utilizing pcDNA3.1 vector with an IRES-GFP. We then performed whole-cell patch-clamp recording of CHO cells expressing E protein. Compared with the un-transfected CHO cells, the cells transfected with SARS-CoV-2 E gene showed moderate inward and relatively larger outward currents under voltage stimulation (Figure. 1, B and C). Using NaMES, K-gluconate and CsMES solution, we further revealed that E protein channels are permeable to monovalent cations Na^+^, K^+^ and Cs^+^ (Figure. 2 A and B). Substituting Na^+^ with the divalent cation Ca^2+^ in the extracellular solution suppressed the inward currents (Figure. 2C), and using NMDG-Cl in the intracellular solution further eliminated outward currents. These results demonstrated that this channel conducts monovalent cations, but not the divalent cation Ca^2+^ and the anion Cl^-^ (Figure. 2 C and D).

**Figure 2.**
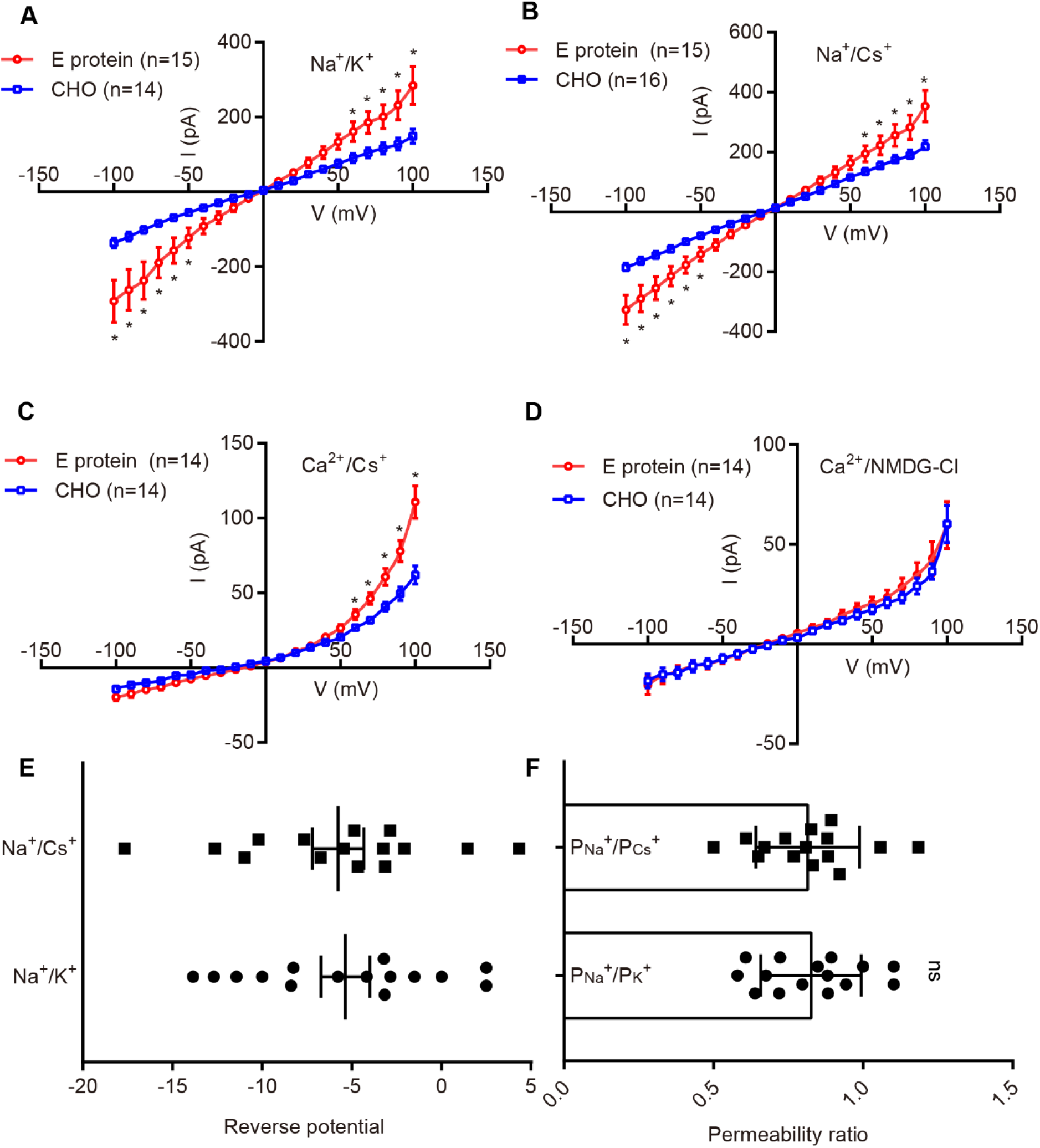
Currents of SARS-CoV-2 E protein in different bi-ionic conditions. **(A)** Currents recorded in extracellular NaMES /intracellular K-gluconate solutions. Data are mean ± SEM. Statistical analysis used T-test; *, p<0.05. **(B)** Currents recorded in extracellular NaMES/intracellular CsMES solutions. Data are mean ± SEM. Statistical analysis used T-test; *, p<0.05. **(C)** Currents recorded in extracellular CaMES /intracellular CsMES solutions. Data are mean ± SEM. Statistical analysis used T-test; *, p<0.05. **(D)** Currents recorded in extracellular CaMES /intracellular NMDG-Cl solutions. Data are mean ± SEM. Statistical analysis used T-test. E protein channels are only permeable to monovalent cations Na^+^, K^+^ and Cs^+^, but not the divalent cation Ca^2+^ and the anion Cl^-^. **(E)** Reverse potentials in different bi-ionic conditions. Data are mean ± SEM. **(F)** Permeability ratios in different bi-ionic conditions. Data are mean ± SD. Statistical analysis used T-test.

We characterized the E protein with reverse potentials of -5.36±1.36 mV (n=15) in Na^+^/ K^+^ solutions and -5.77±1.43 mV (n=15) in Na^+^/ Cs^+^ solutions (Figure. 2 E). The relative permeabilities were calculated as P _Na_ /P _K_ ≈ 0.83 and P _Na_ /P _Cs_ ≈ 0.82 (Figure. 2 F). The numbers did not exhibit significant differences (Figure. 2, E and F).

### SARS-CoV-2 E protein can be inhibited by hexamethylene amiloride

Previous studies demonstrated that the drug hexamethylene amiloride (HMA) blocked the channel activity of SARS-CoV E protein and inhibited virus replication (4,5). Here, we showed that HMA could also inhibit the conductance of the SARS-CoV-2 E protein. With 10 µM HMA application in the bath solution, the channel currents of SARS-CoV-2 E protein were largely reduced to the level of un-transfected CHO cells (Figure. 3). We also demonstrated that the anti-influenza A virus drug amantadine had no effect on SARS-CoV-2 E protein activity with a concentration of 26.6 µM (Figure. S1).

**Figure 3.**
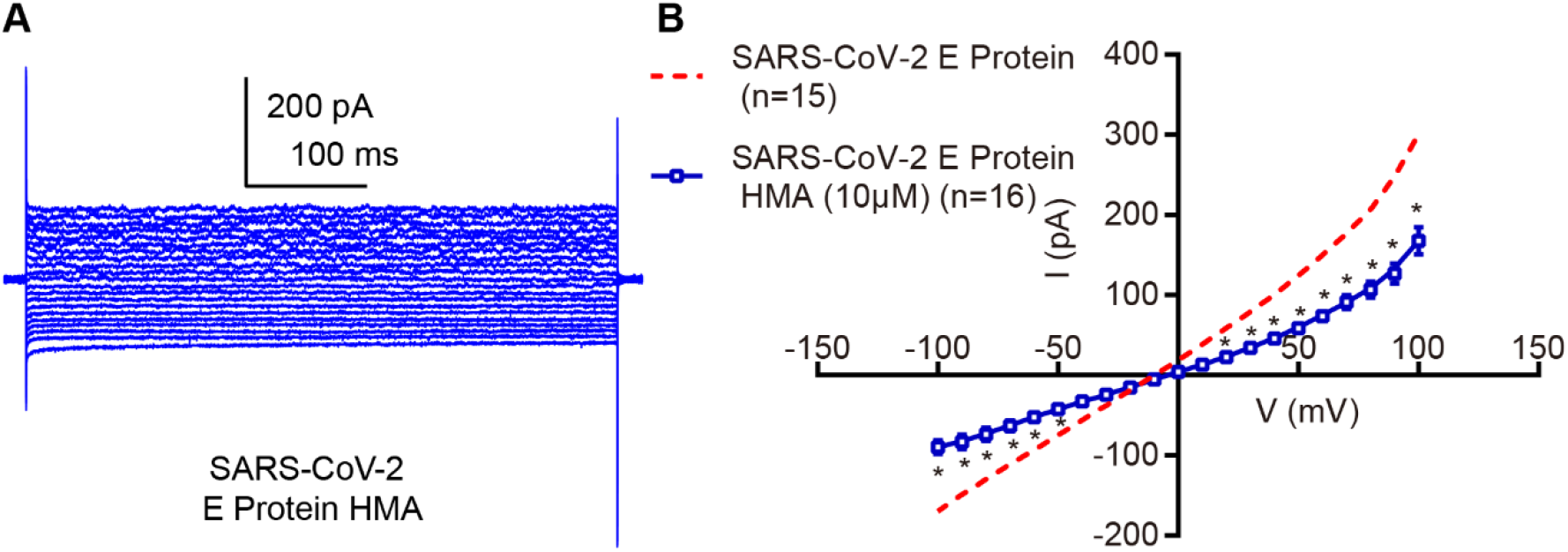
SARS-CoV-2 E protein can be inhibited by HMA. **(A)** Sample traces of CHO cells expressing SARS-CoV-2 E protein with 10 µM HMA. **(B)** Whole-cell current-voltage relationships of the indicated cells. HMA blocked the channel activity of SARS-CoV-2 E protein. The data of E protein channel currents are the same as those in Figure 1C. Data are mean ± SEM. Statistical analysis used T-test; *, p<0.05.

Using homology modeling and the structure of SARS-CoV E protein (6), we built a structural model of the SARS-CoV-2 E protein (Figure. 4A). This model showed a pentameric architecture in which five subunits (A-E) are arranged around a putative central ion permeation path. Each subunit is formed with a N-terminal, a C-terminal and a single transmembrane domain (TMD) (Figure. 4A). Based on the structural model of the SARS-CoV-2 E protein, we further built a binding model of HMA to SARS-CoV-2 E protein. The upper binding pocket was defined by Leu28 in one subunit and Asn45, Ser50, Leu51, Tyr57 in adjacent subunit. The other pocket was composed of Glu8, Glu8, Asn15 in adjacent subunits. By mutating the indicated amino acids in Figure. 4B, we verified the channel activity of the mutants. The results showed that E protein channel currents were abolished in the mutants p.Glu8Ala, p.Glu8Lys, p.Asn15Ala, p.Asn45Ala, p.Ser50Ala and p.Tyr57Ala (Figure. S2), suggesting that these amino acids were critical for the E protein channel activity; consistent with previous studies which showed that p.Asn15Ala mutant abolished SARS-CoV E protein channel activity (2,7). The cells transfected with mutants p.Leu28Ala, p.Leu51Ala produced robust channel currents compared to the un-transfected CHO cells. While wild-type E protein channel was inhibited by HMA (Figure. 3), the currents of mutants p.Leu28Ala, p.Leu51Ala were not suppressed by HMA (Figure. 4). These results showed that Leu28 and Leu51 are critical amino acid mediating HMA inhibition.

**Figure 4.**
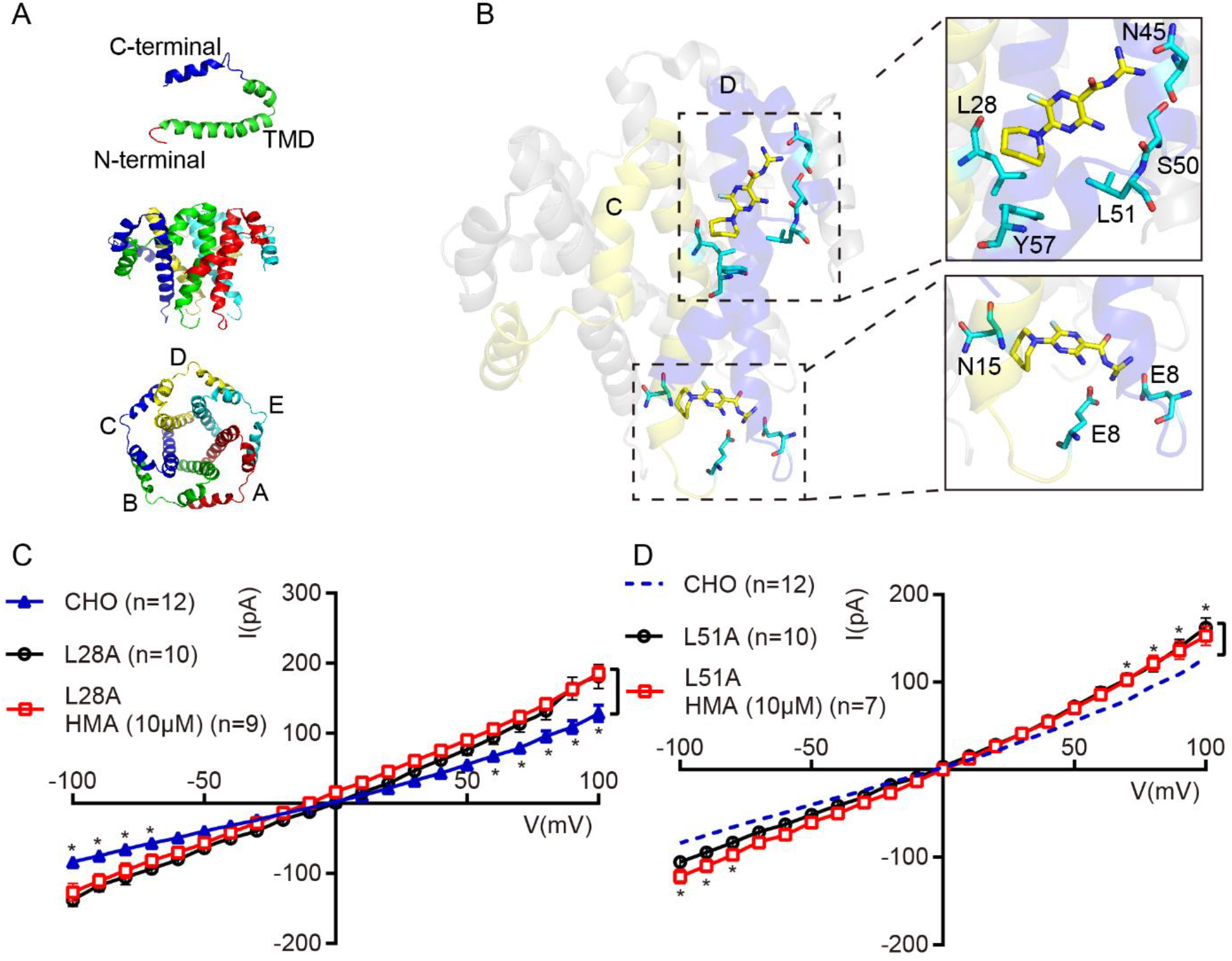
The Binding model for HMA to SARS-CoV-2 E protein. **(A)** The pentameric structural model of SARS-CoV-2 E protein. The five subunits (A-E) are labeled in different colors. **(B)** Binding model of HMA to SARS-CoV-2 E protein. Two binding pockets formed by the interface of adjacent subunits respectively. **(C-D)** Current-voltage relationships of mutants L28A (C), L51A (D) with or without 10 µM HMA. The mutants were not inhibited by HMA and are key determinants for HMA sensitivity. Same data for CHO cell currents in panels C and D. Data are mean ± SEM. Statistical analysis used ANOVA; *, p<0.05.

### Structure-based screening for drugs targeting SARS-CoV-2 E protein

Drug repurposing uses de-risked compounds, thus potentially has lower overall costs and shorter timelines for development (8). Thus, we further used the structural model of SARS-CoV-2 E protein and performed a virtual screening with listed drug library, clinical phase drug library and natural product library with total 5000 compounds (Supplementary table). We obtained 35 candidate compounds (Table 1) and tested their effects on SARS-CoV-2 E protein. We showed that AZD5153, a bromodomain protein 4 inhibitor against hematologic malignancies in clinical trial (9), suppressed the currents of SARS-CoV-2 E protein at 40 µM concentration (Figure. 5A).

**Table 1.** Drug candidates targeting SARS-CoV-2 E protein.

| Approved<br>Status | Clinical Stage | Name | Approved<br>Status | Clinical Stage | Name |
| --- | --- | --- | --- | --- | --- |
| N/A | N/A | 3b-Hydroxy-5-cholenoic<br>acid | N/A | Phase 2<br>Discontinued | L-778123 hydrochloride |
| PMDA | N/A | Temocapil | N/A | N/A | Alprostadil |
| N/A | N/A | 5'-Methylthioadenosine | N/A | N/A | Dinaciclib |
| N/A | Phase 2<br>Discontinued | SCH58261 | N/A | Phase 1<br>Discontinued | SGI-1776 free base |
| N/A | Phase 2<br>Discontinued | Lomeguatrib | N/A | Phase 1<br>Discontinued | PF3274167 |
| N/A | Phase 2 | Vesatolimod | N/A | Phase 2 | Bicalutamide |
| FDA | N/A | Eletriptan HBr | N/A | N/A | 7-Ethylcamptothecin |
| N/A | N/A | Coelenterazine | N/A | Phase 1 | Mivebresib |
| N/A | Phase 1 no<br>development<br>reported | RG2833 | N/A | Phase 2 no<br>development<br>reported | Indometacin |
| N/A | Phase 2 | Quisinostat | FDA | N/A | Azelastine hydrochloride |
| N/A | N/A | Famprofazone | FDA | N/A | Etravirine |
| N/A | Phase 2 | Ganetespib | N/A | Phase 2 | Buparlisib |
| PMDA | N/A | Teneligliptin<br>hydrobromide | N/A | Phase 2<br>Discontinued | ABT639 |
| FDA | N/A | Sulfaphenazole | N/A | Phase 2 | Pleconaril |
| N/A | Phase 2 | ABC294640 | N/A | N/A | Morusin |
| N/A | Phase 1<br>Discontinued | RSV604 | N/A | Phase 1 | AZD5153 6-Hydroxy-2-naphthoic acid |
| FDA | N/A | Rotigotine | FDA | N/A | Droperidol |
| N/A | N/A | Protosappanin B |  |  |  |

**Figure 5.**
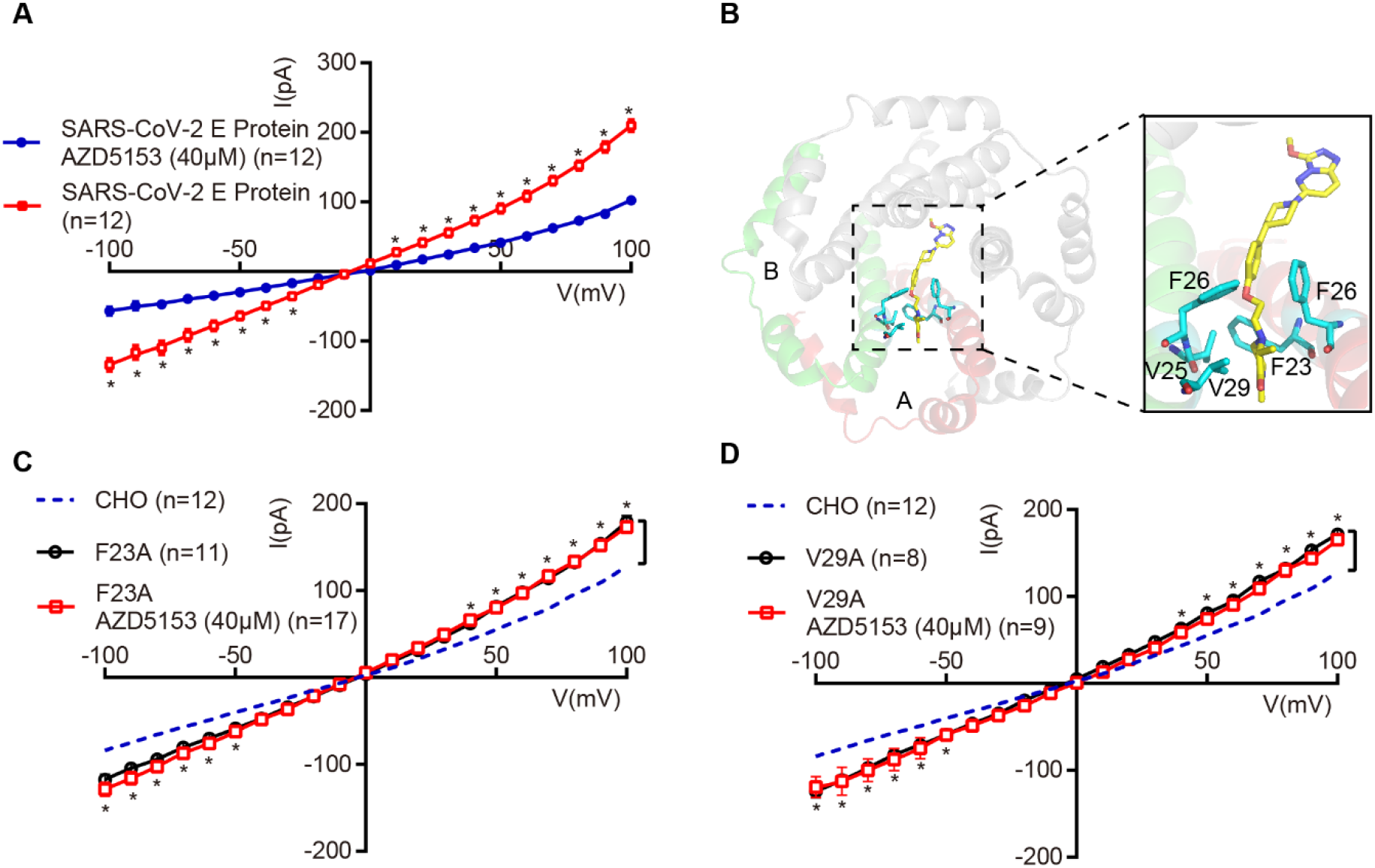
SARS-CoV-2 E protein can be inhibited by AZD5153. **(A)** Whole-cell current-voltage relationships of the indicated cells. SARS-CoV-2 E protein can be inhibited by AZD5153. Statistical analysis used T-test; *, p<0.05. The E protein channel currents recorded varied from that of Figure 1 and Figure 2 due to the use of different batches of cells. (**B)** Binding model of AZD5153 to SARS-CoV-2 E protein. The hydrophobic pocket was formed by the interface of two neighboring subunits. **(C-D)** Current-voltage relationships of mutants F23A (C), V29A (D) with or without 40 µM AZD5153.The mutants lost AZD5153 sensitivity and are crucial for AZD5153’s effects on the SARS-CoV-2 E protein. The data of CHO cell currents are the same as those in Figure 4C. Data are mean ± SEM. Statistical analysis used ANOVA; *, p<0.05.

Figure. 5B showed the binding model of AZD5153 to SARS-CoV-2 E protein. The hydrophobic pocket was formed by the interface composed of Val25, Phe26, Val29 from one subunit, as well as Phe23 and Phe26 from a neighboring subunit. By mutating the indicated amino acids including p.Val25Ala, p. Phe26Ala, p.Val29Ala, p.Phe23Ala, we tested the effect of AZD5153 on the mutants.

We found that all of the mutants still showed channel activity. Further experiments showed mutants p.Phe23Ala and p.Val29Ala lost AZD5153 sensitivity (Figure. 5, C and D) but mutants p.Phe26Ala, p.Val25Ala still can be inhibited (Figure. S3). The results demonstrated that Phe23 and Val29 are key determinants for AZD5153 sensitivity, which is consistent with our binding model prediction.

## DISCUSSION

Our research demonstrated the ion channel activity of SARS-CoV-2 E protein and identified HMA and AZD5153 as its inhibitors. Our point mutation and recording experiments further identified that Leu28, Leu51 are key determinants of HMA inhibition and Phe23, Val29 are required for AZD5153 sensitivity. In Figure.4, we showed that HMA did not inhibit the channel activity of mutants p.Leu28Ala, p.Leu51Ala. The two residues that are critical for E protein and HMA are dominated by non-polar effects. The results strongly supported that HMA binds to the upper pocket consisted of Leu28 in one subunit and Asn45, Ser50, Leu51, Tyr57 in adjacent subunit (Figure.4). In Figure. S2, we showed that all three mutants of another predictive HMA binding pocket (p.Glu8Ala, p.Glu8Lys, p.Asn15Ala) induced E protein channel function loss. In previous studies, the mutation p.Asn15Ala was demonstrated to knock down the channel conductivity of SARS-CoV E protein (10) and further attenuate SARS-CoV virulence (2). In the SARS-CoV E protein, the residues Asn15 and Glu8 were predicted to face the lumen of the channel pore (7,10). In our hypothetical model, HMA might interact with Asn15 and Glu8 (Figure. 4). Given the high homology between the SARS-CoV E protein and SARS-CoV-2 E protein, we speculated that HMA probably binds to a pocket that locates in the pore and thus blocks the channel. A previous study suggested two binding sites of HMA on the SARS-CoV E protein, one in the C-terminus near Arg38 and another in the N-terminus near Asn15. The authors presumed that HMA may bind to the channel by hydrogen bonding to the side-chain carbonyl of Asn15 and the guanidinium moiety of Arg38 (5). The binding sites model near Asn15 in this report was similar to the binding pocket of HMA in SARS-CoV-2 E protein defined by Glu8, Glu8, Asn15 in adjacent subunits in our study.

In Figure. 5, we showed that the residues Phe23 and Val29 were essential for AZD5153 sensitivity. In our predictive binding model, Phe23 and Val29 were two nonpolar amino acids located in two adjacent subunits, which might form non-polar interactions with AZD5153. The SARS-CoV-2 E protein pentamer is composed of five identical monomers. So this binding pattern could be between the A and B subunits, or between the B and C subunits, or between the other two subunits. And it’s possible that multiple compounds could simultaneously bind to the E protein pentamer.

It was reported that the IC_50_ of AZD5153 for BRD4 is <10 nM (9) and therefore this compound is much less potent against the E protein and is unlikely to be an actual therapeutic candidate. It might serve as an interesting starting point for developing more potent inhibitors. A recent research showed that SARS-CoV-2 E protein could cause acute respiratory distress syndrome (ARDS) damages in lungs of mice and inhibitors of SARS-CoV-2 E protein could reduce the viral load in lungs of SARS-CoV-2-infected mice, which suggested that SARS-CoV-2 E protein could be a drug target (11).

In summary, first, our results suggest HMA and AZD5153 as lead compounds to develop other compounds to inhibit SARS-CoV-2 E protein. Second, the established functional essay of SARS-CoV-2 E protein by patch-clamp recording in cultured cells offers an opportunity for drug repurposing such as screening FDA approved drugs as potential drug candidates to treat COVID-19. With this functional essay, we discovered an active molecule against hematologic malignancies as the inhibitor of SARS-CoV-2 E protein.

## EXPERIMENTAL PROCEDURES

### Plasmid construct

The SARS-CoV-2 envelope (E) protein gene (NCBI Reference Sequence: NC_045512.2) was synthesized and subcloned to a pcDNA3.1 vector with an IRES-GFP by GENEWIZ. The E SARS-CoV-2 protein gene variants were generated from the wild-type SARS-CoV-2 E protein gene by homologous recombination method and verified by automated sequencing.

### Cell culture and transfection

Chinese hamster ovary (CHO) cell line was cultured in F-12/DMEM media with 10% fetal bovine serum (FBS) and 1% antibiotic-antimycotic mixture (Invitrogen) at 37 °C with 5% CO_2_. Plasmid was transiently transfected into cells, at a total amount of 2000 ng per dish in 35 mm culture dishes, using Lipofectamine 3000 (Invitrogen) according to the manufacturer’s instructions.

### Electrophysiological recording

Whole-cell voltage-clamp recordings were performed using an Axopatch 700B amplifier (Axon Instruments, USA) at room temperature (20-24 °C). The patch clamp recording was performed 24h after transfection. For patch pipette preparation, the borosilicate glass pipettes (BF150-86-10, Sutter Instrument, Novato) were pulled using a Flaming/Brown micropipette puller (P-97, Sutter Instrument, Novato) and polished to a diameter corresponding to a pipette resistance in the range of 3-6 MΩ. The pipette solution contained (in mM): 135.0 potassium gluconate, 10.0 KCl, 10.0 HEPES, 0.5 EGTA, 2.0 Mg-ATP (pH 7.3 adjusted with KOH; osmolality 276 mOsmkg^−1^) (5). The bathing solution contained (in mM): 124.0 NaCl, 3.5 KCl, 1.0 NaH_2_PO4, 26.2 NaHCO_3_, 1.3 MgSO_4_, 2.5 CaCl_2_ and 10.0 D (+)-glucose (pH 7.4 adjusted with NaOH; osmolarity 300 mOsmkg^−1^).(5) Hexamethylene amiloride (HMA) (A9561, Sigma) was applied in the bath solution at 10 µM concentration. Amantadine hydrochloride (A1260, Sigma) was applied in the bath solution at 26.6 µM concentration. The candidate compounds were purchased from Selleck and applied in the bath solution at 40 µM concentration. Stock solution of HMA at 100mM was prepared in 50% DMSO:50% methanol. Stock solution of amantadine hydrochloride at 25mg/mL was prepared in water. Stock solution of the candidate compounds at 100mM were prepared in DMSO. For ion selectivity experiments, the Cs^+^ pipette solution contained (in mM): 140 CsMES, 10 HEPES (pH 7.2, osmolarity 275 mOsmkg^−1^). The Ca^2+^ bath solution contained (in mM):70 CaMES, 10 HEPES, 100 sucrose (pH 7.2, osmolarity 312 mOsmkg^−1^). The K^+^ pipette solution contained (in mM):140 potassium gluconate, 10 HEPES (pH 7.2, osmolarity 266 mOsmkg^−1^). The Na^+^ bath solution contained (in mM):140 NaMES, 10 HEPES (pH 7.2, osmolarity 280 mOsmkg^−1^) (12). The NMDG pipette solution contained (in mM):140 NMDG, 10 HEPES (pH 7.4 adjusted with HCl, osmolarity 263 mOsmkg^−1^). Signals were filtered at 1 kHz (low-pass) and digitized using a DigiData 1550 with pClamp 10.5 software (Molecular Devices, USA). The sampling rate was 5 kHz. The pipette off-set was corrected after the pipette entered the bath solution. C-fast cancellations were done once a GΩ seal has been formed. Whole-cell capacitance compensation was performed after the cell membrane was broken. The serial resistance was monitored throughout the experiments.

Series resistance compensation (>80%) was performed in the experiments. The whole-cell current traces of different groups were generated by voltage steps ranging from -100 to 100 mV with 10 mV increments from a holding potential of 0 mV. The currents at different voltage potentials were measured at peak levels to generate the current-voltage relationship. Analyses of data were performed with GraphPad Prism7. Pooled data are shown as means±SEM. Data were analyzed by unpaired *t* test or ANOVA.

### Determination of permeability ratios

For determination of the relative permeability ratio of P_X_/P_Y_ for monovalent cations, we used the simplified Goldman-Hodgkin-Katz (GHK) equation:

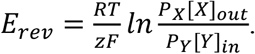

At 23°C, RT/zF has the value of 25.5. E_rev_ represents the reversal potential of the current. Reversal potential for each cell in different bi-ionic condition was calculated from interpolation of the relative current-voltage data. Reversal potentials and permeability ratios were presented as mean ± SEM.

### Structure modeling

Modeling of SARS-CoV-2 E protein was carried out on SWISS-MODEL server. The protein sequence of SARS-CoV-2 E protein (YP_009724392.1) was obtained from NCBI database. PDB structure of SARS-CoV E protein (PDB ID: 5×29), which shares homology of 94.7% with SARS-CoV-2 E protein, was used as a template. Furthermore, a QMEAN score of -9.66 confirmed the reliability of the homology structural model of SARS-CoV-2 E protein.

### Virtual screening

The SiteMap protocol was used to predict the possible binding pocket of SARS-CoV-2 E protein in Schrodinger. As a result, three possible binding pocket was identified. The pocket located within the possible central ion permeation path was chosen to perform the virtual screening.

Structural optimization was conducted in Schrodinger with the default protocol of the Protein Preparation Wizard. Energy minimization was carried out using the OPLS-2005 force-field. Since the five monomers of SARS-CoV-2 E are identical in the pentameric model, amino acids 12-27 of monomer A and B was chosen as a binding pocket to build a docking grid using the Receptor Grid Generation tool. The box dimensions for the grid box was 20 Å × 30 Å × 20 Å. The virtual screening used the listed drug library, clinical phase drug library and natural product library in Targetmol (Supplementary table). The QikProp module was used for efficient evaluation of pharmaceutically relevant properties of Targetmol compounds library; it predicts the Absorption, Distribution, Metabolism, Elimination (ADME) properties of all compounds. Lipinski’s Rule of Five was used to filter out the unqualified small molecules. All small compounds were prepared using the MMFFs force-field in the LigPrep module.

High throughput virtual screening (HTVS) was carried out by Schrodinger. We have screened the L6000 Targetmol compound library against the SARS-CoV-2 E protein structure. Compounds which were screened successfully from HTVS were further subjected to SP (standard-precision) docking to get more accurate results. Furthermore, XP (extra precision) docking was used to remove the false-positive results.

The top-ranked 283 molecules obtained from this virtual screening were then grouped into hierarchical clusters based on their chemical similarity calculated using Tanimoto similarity scores of binary fingerprints in the Canvas. From each cluster, the small compounds with the highest docking score were selected to retain, and finally 35 small compounds with different chemical structures were obtained.

## DATA AVAILABILITY

All data are contained within the manuscript.

## SUPPORTING INFORMATION

This article contains supporting information.

## ACKNOWLEDGMENTS

Thanks Woo-Ping Ge, Jun Li and Weizhi Sun for discussion.

## AUTHOR CONTRIBUTIONS

X.X: investigation, formal analysis, methodology, visualization, writing – original draft, writing – review & editing; Y. Z: investigation, formal analysis, visualization, writing – original draft;S. L: investigation, formal analysis, visualization, writing – original draft; H. L: investigation, formal analysis; Z. Y: conceptualization literature search, funding acquisition, project administration, supervision, writing – original draft, writing – review & editing.

## FUNDING AND ADDITIONAL INFORMATION

The research was supported by funds from the National Key Research and Development Program of China Stem Cell and Translational Research (2017YFA0103900), the National Natural Science Foundation of China (31571083, 31970931), the Shanghai Municipal Science and Technology Major Project (No.2017SHZDZX01 and No.2018SHZDZX01) and ZJLab.

## CONFLICT OF INTERESTS

The authors declare that they have no conflicts of interest with the contents of this article.

## Notes

### Competing Interest Statement

The authors have declared no competing interest.

