## Supplementary figures for "Structure-based screening of drug candidates targeting the SARS-CoV-2 envelope protein"

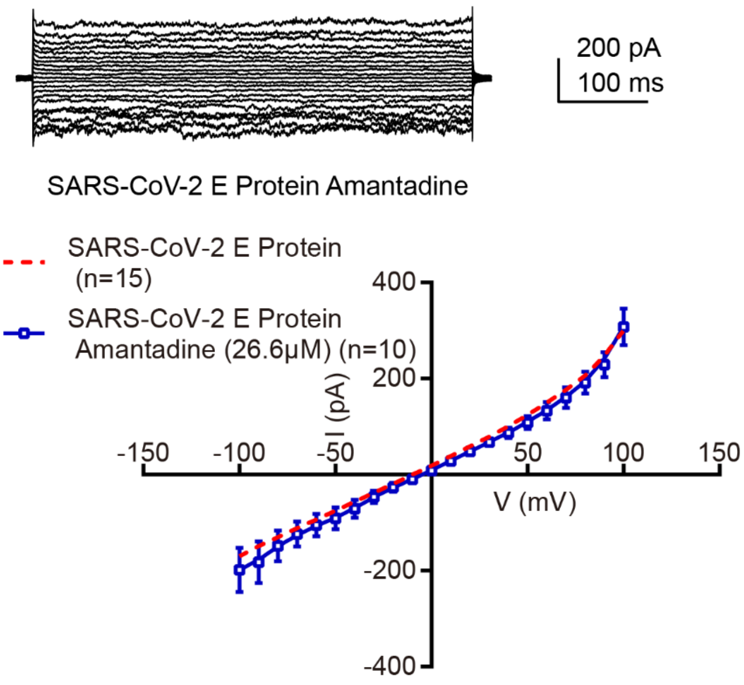


**Figure S1.** Currents of SARS-CoV-2 E protein with the application of amantadine. The current traces were generated using the same protocol in Figure 1. Amantadine was applied in the bath solution at the concentration of 26.6 μM and had no effect on SARS-CoV-2 E protein activity. The data of E protein channel currents are the same as those in Figure 1C. Statistical analysis used T-test.

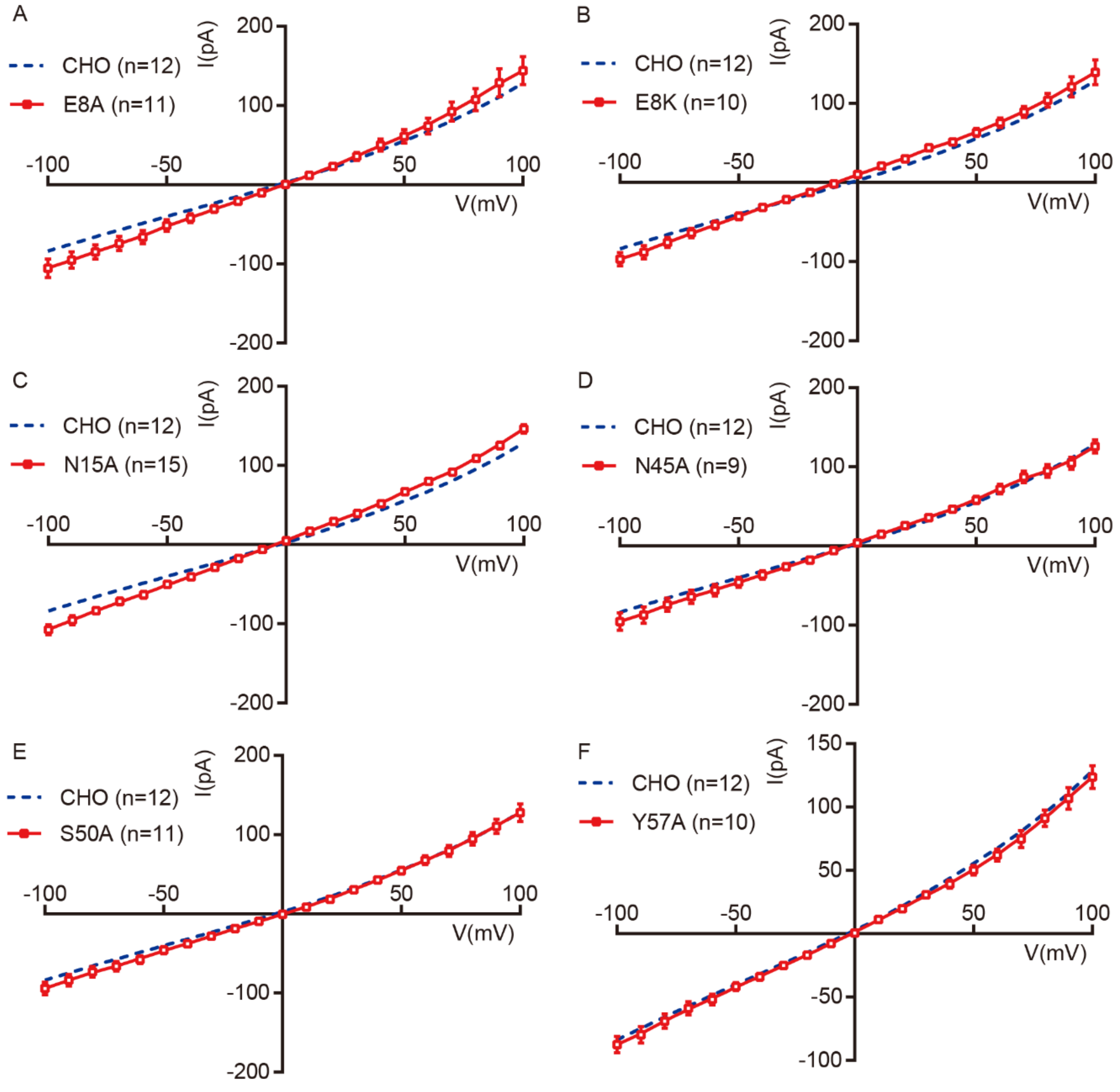


**Figure S2.** Currents of SARS-CoV-2 E protein mutants. Current-voltage relationships of un-transfected CHO cells and mutants E8A (A), E8K (B), N15A(C), N45A(D), S50A (E), Y57A(F). The mutants did not show channel activity. The data of CHO cell currents are the same as those in Figure 4C. Data are mean ± SEM. Statistical analysis used T-test.

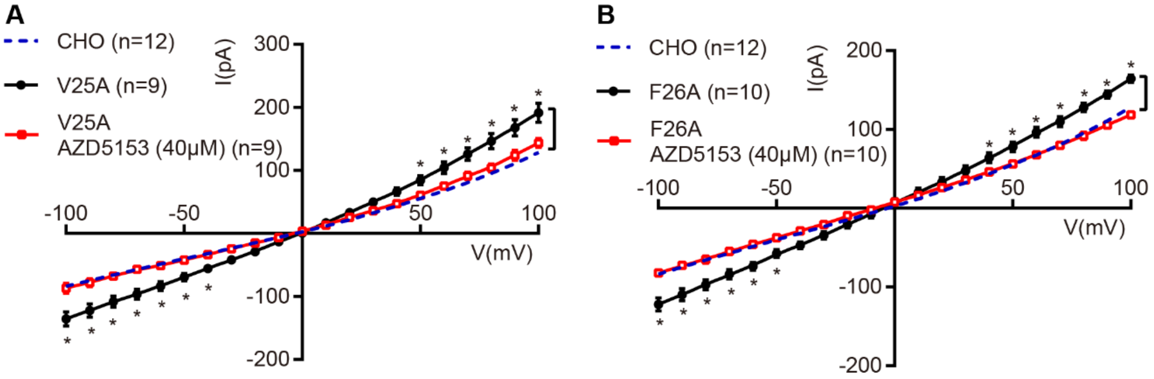


**Figure S3.** Currents of SARS-CoV-2 E protein mutants with or without AZD5153. Current-voltage relationships of un-transfected CHO cells and mutants V25A (A), F26A (B) with or without 40 μM AZD5153. The currents of un-transfected CHO cells showed differences compared with that of the mutants transfected cells. The mutants can be inhibited by AZD5153. The data of CHO cell currents are the same as those in Figure 4C. Data are mean ± SEM. Statistical analysis used ANOVA; *, $p<0.05$.
